# Experimental characterization of zeta viruses and discovery of betaviroid-like RNAs expand the diversity of coding circular RNA replicons

**DOI:** 10.64898/2026.09.16.752053

**Authors:** Nadia Serale, Michela Chiumenti, Paolo Mussano, Silvia Rotunno, Francesco Venice, Gianni Della Rocca, Massimo Turina, Antonietta Mello, Laura Miozzi, Francesco Di Serio, Beatriz Navarro

**Author notes:** Address correspondence to Beatriz Navarro and Francesco Di Serio.

## Abstract

Viroid-like RNAs (vdlRNAs) are infectious agents characterized by single-stranded circular genomes folding into compact secondary structures, often containing self-cleaving ribozymes involved in replication. Recent metatranscriptomic studies have expanded their diversity across multiple ecological niches and revealed novel vdlRNAs groups with protein-coding capacity, including zeta viruses, characterized by the potential to encode an endless tandem-repeat protein in each polarity strand. However, besides their *in silico* identification, no experimental evidence supported their existence and molecular features. Here, three new zeta viruses were identified and one of them was molecularly characterized from soil. In addition, a novel type of coding vdlRNAs, named betaviroid-like RNAs was discovered. Similar to zeta viruses, betaviroid-like RNAs are circular RNAs containing, in each polarity strand, one ribozyme and one open reading frame (ORF) encompassing the full-length genome. Unlike zeta viruses, betaviroid-like ORFs are not endless but longer than the unit-length genome. Existence of both circular RNA polarity strands, absence of DNA forms, and self-cleavage activity of ribozymes were experimentally confirmed for both zeta viruses and betaviroid-like RNAs, providing data consistent with symmetric rolling-circle replication. Additional zeta viruses and betaviroid-like RNAs were identified in publicly available metatranscriptomic datasets from wetlands of different geographic areas, supporting their broad distribution. Sixteen zeta viruses contain diverse self-cleaving ribozymes, including delta, type II and III hammerhead, twister, and hairpin ribozymes, expanding the known ribozyme diversity of this group. Altogether, our results provide the first experimental characterization of zeta viruses and betaviroid-like RNAs expanding the diversity of circular RNA replicons.

**IMPORTANCE:** This study supports the existence of two novel groups of protein-coding viroid-like RNAs, further expanding the known diversity and complexity of these agents. In addition to exhibiting the characteristic features of viroid-like RNAs, including circular RNA genomes, compact secondary structures, and self-cleaving ribozymes active in both polarity strands, these molecules possess coding capacity on both strands, with the coding potential exceeding the length of the genomic RNA itself. The coexistence of replication-associated RNA motifs and protein-coding sequences within the same genomic regions points to an extraordinary level of genomic information compression and reveals a previously unrecognized strategy of genome organization. At the same time, these findings raise important questions regarding the origin, evolution, expression, and biological roles of these RNAs, as well as the selective pressures that have shaped such a highly compact genomic architecture.

## INTRODUCTION

Viroid-like RNAs (vdlRNAs) are a group of infectious agents characterized by a single-stranded circular RNA genome that adopts a highly compact, base-paired secondary structure. Replication of vdlRNAs generally occurs through a rolling-circle replication mechanism that proceeds via RNA intermediates and is mediated by host enzymes. In several cases, they have self-cleaving ribozymes, which catalyze the precise cleavage of RNA replication intermediates involved in their replication. Depending on the group, vdlRNAs may be non-coding or contain one or more open reading frames (ORFs), that code for proteins with putative roles in replication or other aspects of their life cycle (1). This broad category includes multiple agents that differ in genome size, host range, and replication strategy. Among viroid-like RNAs, viroids are the smallest infectious agents [approximately 200–450 nucleotides (nt)] reported so far. Consisting of circular non-coding RNAs, they replicate autonomously and infect plants systemically (2). Viroid-like satellite RNAs structurally resemble viroids and also infect plants, but depend on a helper virus for infectivity (3). Hepatitis delta virus (HDV) (4) and deltavirus-like RNAs (5) possess larger circular genomes (approximately 1,200–2,000 nt) than viroids and viroid-like satellite RNAs, encode the delta antigen or delta-like proteins, and infect humans and other animals. Another group of viroid-like elements are retrozymes and retroviroids, which however are not infectious and have a DNA counterpart integrated in animal or plant host genome (6–9). Recently, metatranscriptomic analyses have revealed that vdlRNAs are more widespread than previously thought and occur in several ecological niches, with host range extended to fungi and bacteria (10–14).

A distinct group of vdlRNAs recently described is represented by zeta viruses, which have been identified exclusively *in silico* so far (15). To date, approximately 311 zeta virus sequences have been detected in metatranscriptomic datasets from environmental and invertebrate-associated samples, although their actual hosts remain unknown and no experimental evidence of their existence has been provided so far (15). Zeta viruses are reported to be circular RNAs (324–789 nt) and are predicted to fold into rod-like secondary structures. A defining feature of this group is that their genomic RNA is composed of a number of nucleotides that is multiple of three and encodes two ORFs, one in each polarity strand, each spanning the entire genome and lacking stop codons. This unusual feature suggests the potential production of endless tandem repeats of a zeta virus protein from both genomic polarities. In addition, both polarity strands contain self-cleaving hammerhead ribozymes (15).

Here, data providing the first experimental identification and molecular characterization of a zeta virus are presented. Moreover, we describe a novel group of coding vdlRNAs, termed betaviroid-like RNAs, characterized by the presence of both a ribozyme and an ORF in each polarity strand. Notably, the ORF lacks an in-frame termination codon within the monomeric RNA molecule. Instead, translation termination is generated only after a frameshift event in a putative multimeric RNA replication intermediate, or through reiterative translation of the circular RNA. In both scenarios, the resulting coding sequence extends beyond the length of the monomeric vdlRNA genome. Data supporting the occurrence of betaviroid-like RNAs in nature are also shown and discussed.

## RESULTS

### Identification and characterization of new zeta viruses

Searches performed by INFERNAL algorithm (16) for the identification of RNAs containing ribozymes showed, in three out of the six RNAseq libraries (C30, C105, and C110) from soil, 109, 32, and 17 contigs, respectively, corresponding to putative vdlRNAs. Among these contigs, we selected three that exhibited the characteristic features of zeta viruses for further characterization. The contigs NODE_21658 (647 nt length; 3.67 coverage) and NODE_1090 (721 nt length; 11.18 coverage) from C105 and C110 libraries, respectively, correspond to the same putative circular RNA of 600 nt, named ZV1 (Fig. 1A). The contigs NODE_198789 (591 nt length; 22.62 coverage) and NODE_416260 (447 nt length; 0.52 coverage), both from C30 library, correspond to two putative circular RNAs of 405 and 399 nt, named ZV2 and ZV3, respectively (Fig. 1A). These RNAs have a nucleotide sequence identity with each other between 52 and 58.5% and, based on BLAST searches, did not display any significant identity with other previously reported sequences. A type III hammerhead ribozyme (HHRz) was contained in each polarity strand of the three RNAs (Fig. 1B), with HHRz sequences paired in the predicted rod-like secondary structure (Fig. 1A). One HHRz of ZV1, located in the (-) strand according with data reported later, shows a mutation in the conserved catalytic core (the typical motif CUGA is replaced by CCGA). All these vdlRNAs contain a single open reading frame (ORF) on each polarity strand, with each ORF spanning the entire (circular) genome without an in-frame stop codon. As a result, each strand of ZV1, ZV2 and ZV3 putatively encode a protein consisting of an endless tandem repeat of a sequence unit consisting of 200, 135 and 133 amino acids (aa), respectively. No additional ORFs were identified when searches were performed using genetic codes other than eukaryotic one. The compact circular, rod-like RNA secondary structure, the presence of paired HHRzs, and a genome length that is a multiple of three (405, 399, and 600 nt), which enables uninterrupted translation of the circular RNA into putative endless tandem-repeat proteins, constitute the defining features of zeta viruses.

**Fig. 1.**
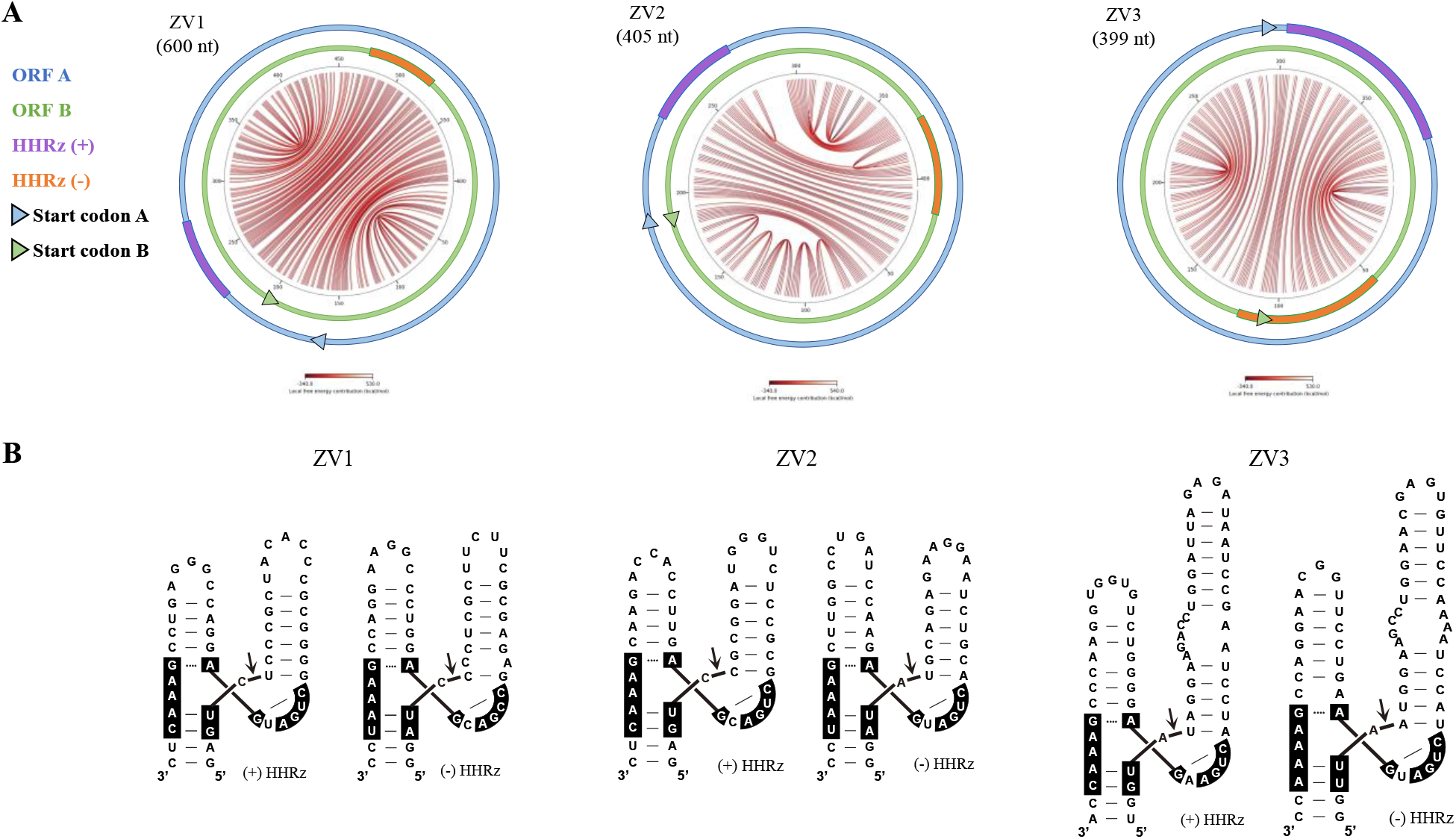
Structural features of zeta viruses. (A) Circular plot representation of the minimum free energy secondary structure of ZV1, ZV2, and ZV3 RNAs. Predicted base-pairing interactions within the RNA genome are indicated by arcs, whose colours, ranging from white to dark red, reflect base-pair stability expressed as local free energy contribution (kcal/mol). The relative positions of (+) and (-) hammerhead ribozymes (HHRz) are indicated in purple and orange, respectively. ORF A [(+) strand] and ORF B [(-) strand] are shown in blue and green, respectively, with triangles marking the corresponding start codons. (B) Secondary structure of (+) (on the left) and (-) HHRzs (on the right) of ZV1, ZV2, and ZV3. The cleavage site is marked by an arrow and conserved nucleotides within the catalytic core are highlighted with a black background.

The six putative monomeric zeta virus proteins shared low aa sequence identity with each other, ranging from 11.3% to 24.8%. The tertiary structures of the monomeric zeta virus proteins, predicted by AlphaFold 3, showed predominantly α-helix motifs (Fig. S1). However, the associated template modelling (pTM) scores were low, ranging from 0.16 to 0.24, indicating limited confidence in the structural models (pTM < 0.5). Predicted structures of the putative dimeric and trimeric proteins, potentially generated by reiterative translation of circular or multimeric zeta virus RNAs, likewise displayed predominantly α-helix motifs, but these models were also associated with low pTM scores (Fig. S1).

HTS sequencing data were analyzed to assess the sequence variability in zeta virus genomes. Sequence variability was expected to be highly constrained because the genome must simultaneously preserve conserved RNA structural features, including the compact secondary structure and self-cleaving ribozymes, while maintaining coding capacity on both polarity strands. Furthermore, the potential ability to encode tandem-repeat proteins requires the genome length to remain a multiple of three nucleotides. A total of 150 and 627 reads from C105 and C110 libraries, respectively, mapped within the ZV1 sequence. Analysis of the aligned reads revealed a single polymorphic site (G→C s at position 137) in the C110 library, with a frequency of 57.1% (variant p-value = 6.3 × 10⁻⁵). This nucleotide substitution did not introduce premature stop codons in either of the putative proteins encoded by the two ORFs. However, it resulted in a non-conservative amino acid substitution (Tryptophan → Serine) in the potential protein encoded by ORF A and a conservative amino acid substitution (Glutamine → Glutamic acid) in the protein encoded by ORF B. In the C30 library a total of 508 reads mapped within the ZV2 sequence, allowing the identification of two polymorphic sites (G→A and T→A substitutions at position 59 and 16, respectively). These variants occurred at frequencies of 69.1% (variant p-value = 6.8 × 10⁻^130^) and 35.6% (variant p-value = 4.7 × 10⁻^92^), respectively. Neither substitution generated premature stop codons in the putative proteins encoded by the two ORFs. The G→A mutation resulted in non-conservative amino acid substitutions in both predicted proteins, causing Glycine → Glutamic acid and Proline → Serine changes in the products of ORF A and ORF B, respectively. Likewise, the T→A substitution resulted in non-conservative amino acid replacements in both putative proteins, namely Tryptophan → Arginine in ORF A and Glutamine → Leucine in ORF B. Only eight reads from library C30 mapped to the ZV3 sequence, and no polymorphic sites were detected, most likely because of the very low sequencing depth. Importantly, no nucleotide insertions or deletions were identified in any of the putative vdlRNAs analysed, indicating strong conservation of genome length and reading-frame integrity.

For further characterization, we used the ZV1 as model. To detect ZV1, RNA preparations from soil samples were reverse transcribed using random primers, followed by PCR amplification with two adjacent specific primers of opposite polarity (ZetaV_1 and ZetaV_2, Table S1) able to amplify circular monomeric and/or multimeric forms. An amplicon with the size consistent with the full-length genome of ZV1 was obtained from both soil samples C105 and C110 (Fig. 2A). Cloning and Sanger sequencing of the amplified cDNAs confirmed that their sequence corresponded to the ZV1 genomic RNA identified *in silico*, providing experimental evidence of its existence (ZV1 GeneBank accession number: PZ584207).

**Fig. 2.**
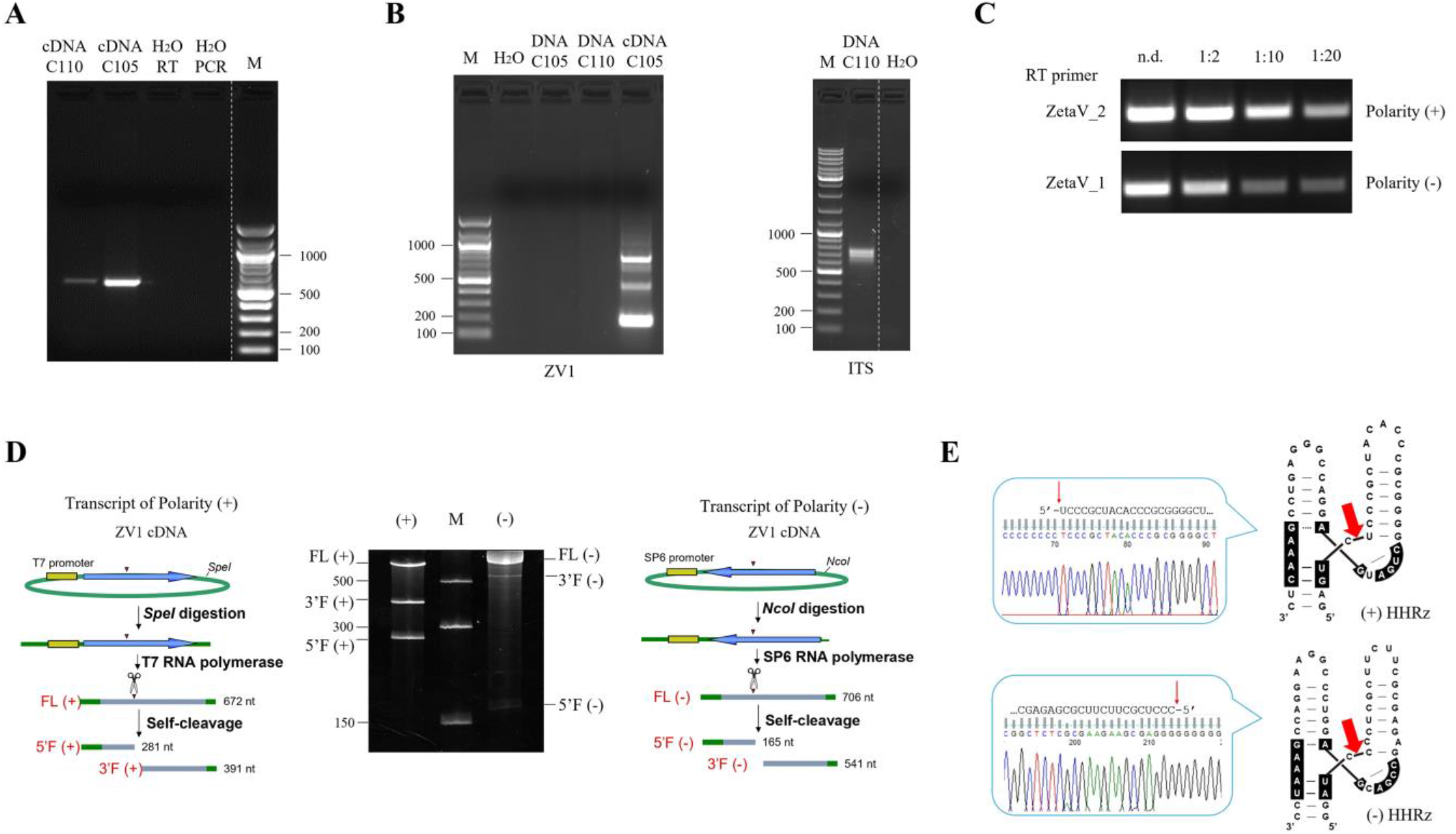
Molecular characterization of zeta virus 1 (ZV1). (A) RT-PCR amplification of genomic full-length ZV1, using adjacent primers of opposite polarity, from C105 and C110 soil samples. (B) PCR amplification of a ZV1 fragment (184 nt) using specific primers (ZetaV_184_3 and ZetaV_184_4) and either cDNA or DNA preparations as template, to confirm the absence of DNA counterpart (on the left). PCR amplification of DNA preparations using fungal universal primers ITS4/ITS5 were used as positive control (on the right). (C) Strand-specific RT followed by semi-quantitative PCR for the determination of the ZV1 most abundant polarity strand denoted as (+) polarity by convention. n.d., not diluted. M, Gene ruler DNA ladder mix (Thermo Scientific). (D) Schematic representation of the DNA templates and monomeric RNA products generated by *in vitro* transcription. Plasmid containing the monomeric ZV1 sequence linearized with SpeI and transcribed with T7 RNA polymerase generates monomeric transcripts (Full length, FL) of positive polarity; the size of the expected 5’ and 3’ fragments (5’F and 3’F) derived from the HHRz self-cleaving activity are indicated (left side). Linearization with NcoI and transcription with SP6 RNA polymerase of the same plasmid produce monomeric transcripts (FL) of negative polarity and their corresponding self-cleavage 5’F and 3’F fragments (on the right). In green, plasmid sequences; in yellow, polymerase promoter; in blue, ZV1 sequence, with the arrows indicating the orientation; arrowheads and scissors mark the position of the self-cleavage sites. Expected sizes of the RNA fragments are indicated on the left. The middle panel shows PAGE analysis of *in vitro* transcription reaction of positive and negative ZV1 monomers. M, RNA marker (Low range ssRNA ladder, New England Biolabs), with sizes (nt) indicated on the left. (E) Determination of self-cleavage site (red arrow) by 5′ RACE of the 3′F fragment resulting from HHRz mediated activity. Sequencing electropherograms of 5′ RACE products of the (+) and (−) 3′F self-cleavage fragments are shown on the left, upper and lower panels, respectively. The self-cleavage sites (red arrow) of the (+) and (-) HHRz (on the right) were determined by 5’RACE of the self-cleavage 3’F fragments. Ribozyme secondary structures are shown on the right.

To investigate whether ZV1 RNA has a DNA counterpart, cDNA and DNA preparations from C105 and C110 soil samples were used for PCR amplification with ZV1-specific primers (ZetaV_184_3 and ZetaV_184_4, Table S1), designed to amplify a partial sequence of 184 nt. Amplification of fungal ITS by universal primers ITS4/ITS5 (Table S1) was used as positive control. Indeed, the presence of fungal DNA in these preparations is expected according to previous published data (17). No amplification product was detected with ZV1 specific primers when DNA was used as template, whereas an amplicon of the expected size was obtained using the fungal primers in the same template and using the ZV1 specific primers on cDNA preparations (Fig. 2B). These results indicated that ZV1 lacks a DNA counterpart.

To assess the existence of the two RNA polarities and to assign the polarity (+) to the strand accumulating at higher level, a semi-quantitative RT-PCR was performed. Each polarity strand was separately retro-transcribed using either ZetaV_1 and ZetaV_2 strand-specific primers. Several dilutions (not diluted, 1:2, 1:10, 1:20) of the resulting cDNAs were then amplified by PCR using the primers ZetaV_184_3 and ZetaV_184_4. Both RNA polarities were amplified and a substantially higher steady state level of the ZV1 RNA polarity transcribed with ZetaV_2 was observed (Fig. 2C), which was therefore considered as the (+) polarity strand.

The self-cleavage activity of ZV1 hammerhead ribozymes was assessed by *in vitro* transcription of the monomeric genomic RNA of both polarity strands. The full-length transcript and the two products expected to be generated by the hammerhead-mediated self-cleavage were detected in both cases (Fig. 2D). Moreover, the precise self-cleavage site of the (+) and (-) HHRzs were confirmed by 5’ RACE analysis, which showed that the 3’ fragments generated by the ribozymes activity during transcription had the expected 5’ termini (Fig. 2E). Altogether these data confirmed that the hammerhead ribozymes of ZV1 are active *in vitro*, suggesting a replication through a symmetric rolling circle mechanism.

### Identification of zeta virus RNAs containing self-cleaving ribozymes other than type III hammerhead ribozymes

Hammerhead ribozymes are divided into three topological classes (I, II, and III) according to which helix contains the termini of the RNA molecule (18). To date, the zeta viruses described in literature are characterized by the presence of type III HHRzs in both genomic polarities (15). To investigate the existence of zeta viruses containing other type of self-cleaving ribozymes, a search was conducted on the dataset of viroid-like RNAs previously identified *in silico* by Forgia et al. (10). Interestingly, this search revealed 16 zeta viruses (Fig. 3) containing self-cleaving ribozymes other than type III HHRz in different combinations (Table 1 and Fig. 4). The newly identified zeta viruses exhibited a rod-like or quasi rod-like predicted secondary structure. The only notable exception was the zeta virus ZV130856 which displayed a considerably more branched structure (Fig. 3).

**Fig. 3.**
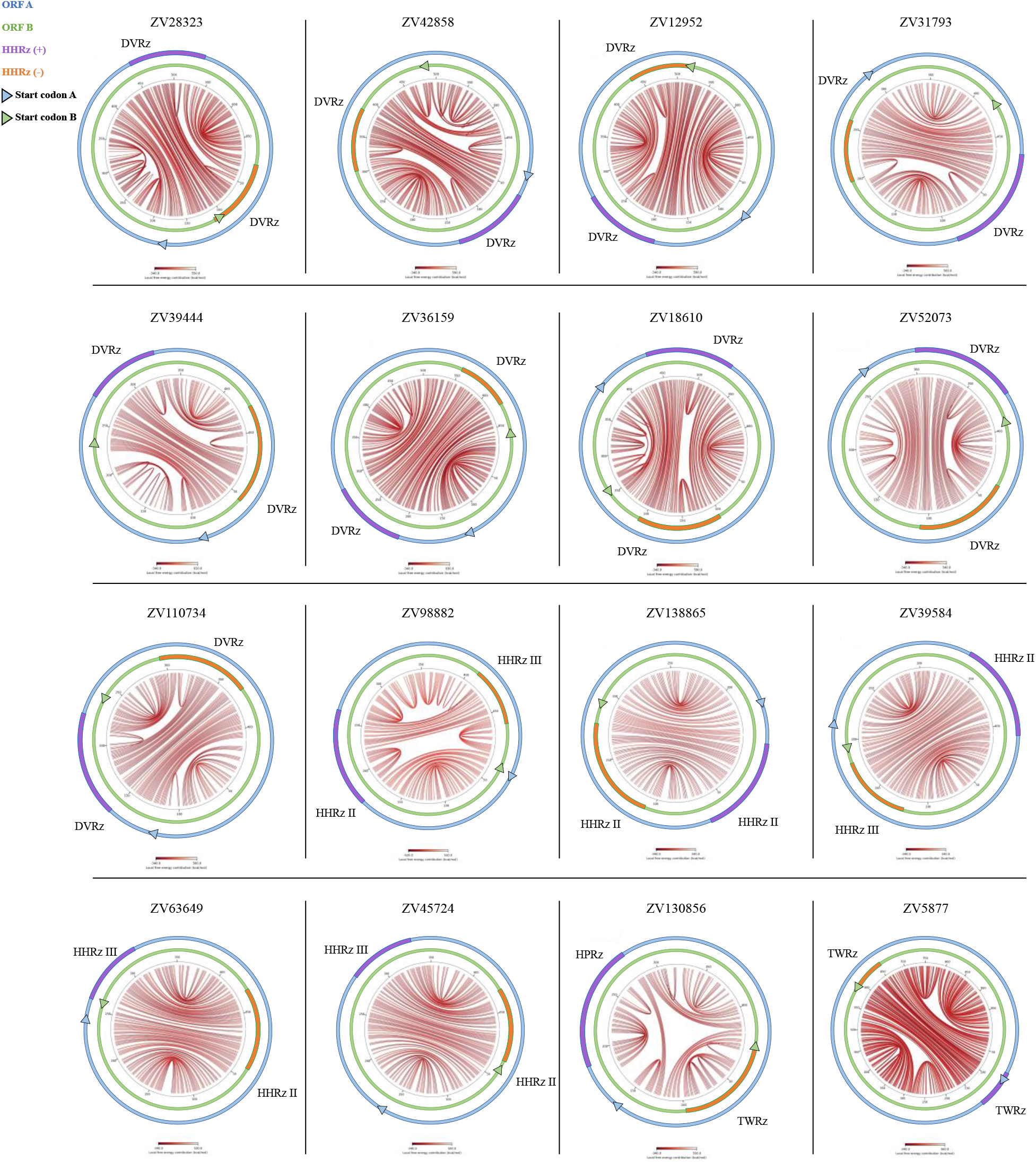
Circular plot of the minimum free energy secondary structure of zeta viruses with ribozymes different than type III HHRz identified in metatranscriptomic datasets. The relative positions of (+) and (-) ribozymes are indicated in purple and orange, respectively. ORFs encoded in (+) and (-) strands are shown in blue and green, respectively, with triangles marking the corresponding start codons (see legend Fig. 1 for details).

**Fig. 4.**
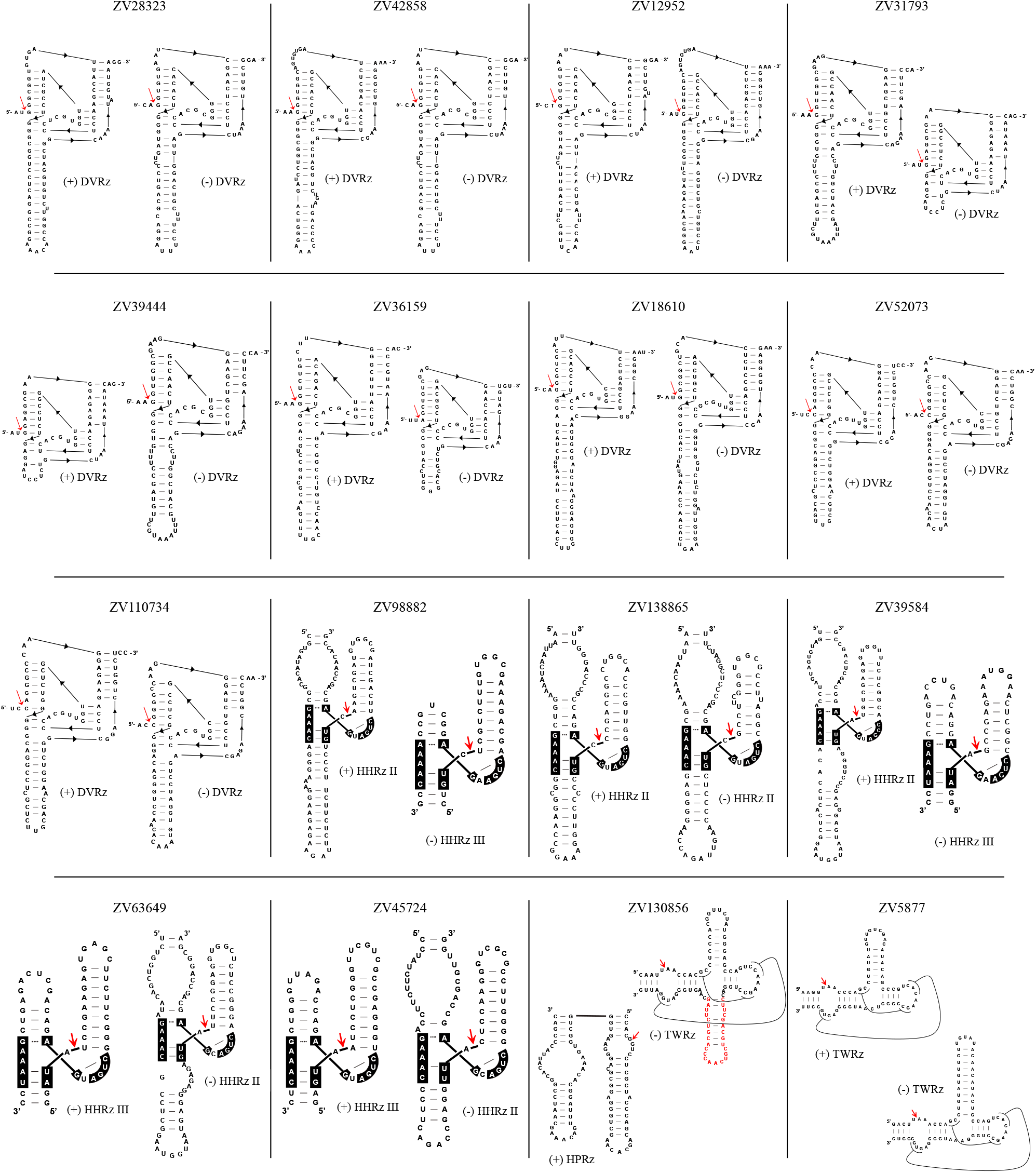
Secondary structure of (+) and (-) ribozymes found in zeta viruses from metatranscriptomic datasets and not containing paired type III HHRz. DVRz: delta ribozyme; HHRz II: type II hammerhead ribozymes; HHRz III: type III hammerhead ribozymes; HPRz: hairpin ribozyme; TWRz: twister ribozyme. The cleavage site is marked by a red arrow and conserved nucleotides within the catalytic core are highlighted with a black background. Nucleotides forming the additional helix-loop element identified in the twister ribozyme in one polarity of ZV130856 are highlighted in red.

**Table 1.** List of putative zeta viruses containing ribozymes other than type III HHRzs.

| ID | Contig | Genome<br>size (nt) | (+)<br>Ribozyme | E-value | (-)<br>Ribozyme | E-value |
| --- | --- | --- | --- | --- | --- | --- |
| **ZV28323** | SRR11829260;NODE_28323_length_742_cov_26.500747 | 669 | DVRz | 5.7e-05 | DVRz | 5.7e-05 |
| **ZV42858** | SRR10482243;NODE_42858_length_739_cov_170.015015 | 666 | DVRz | 8.4e-05 | DVRz | 8.6e-03 |
| **ZV12952** | SRR7428111;NODE_12952_length_715_cov_3.442943 | 666 | DVRz | 1.4e-03 | DVRz | 1.2e-04 |
| **ZV31793** | SRR11450578;NODE_31793_length_535_cov_103.549784 | 462 | DVRz | 8.0e-05 | DVRz | 1.5e-04 |
| **ZV39444** | SRR5864101;NODE_39444_length_535_cov_1.967532_1 | 462 | DVRz | 7.6e-04 | DVRz | 8.0e-05 |
| **ZV36159** | SRR11614418;NODE_36159_length_751_cov_5.014749 | 678 | DVRz | 1.1e-04 | DVRz | 9.0e-03 |
| **ZV18610** | SRR8512518;NODE_18610_length_676_cov_17.376396 | 627 | DVRz | 1.5e-04 | DVRz | 8.1e-02 |
| **ZV52073** | SRR7286070;NODE_52073_length_484_cov_65.367397 | 411 | DVRz | 2.8e-02 | DVRz | 1.9e-04 |
| **ZV110734** | SRR8857868;NODE_110734_length_484_cov_15.513382 | 411 | DVRz | 2.8e-02 | DVRz | 9.4e-03 |
| **ZV130856** | SRR11565845;NODE_130856_length_499_cov_8.208920 | 426 | HPRz | 3.1e-08 | TWRz | 1.2e-06 |
| **ZV98882** | SRR11828805;NODE_98882_length_550_cov_4.985325 | 477 | HHRz II | 3.2e-09 | HHRz III | 2.2e-01 |
| **ZV138865** | SRR11011818;NODE_138865_length_412_cov_3.073746 | 339 | HHRz II | 6.7e-04 | HHRz II | 3.2e-07 |
| **ZV39584** | SRR5839026;NODE_39584_length_481_cov_9.259804 | 408 | HHRz II | 9.3e-04 | HHRz III | 3.9e-08 |
| **ZV63649** | SRR12264489;NODE_63649_length_541_cov_16.369658 | 468 | HHRz III | 1.8e-04 | HHRz II | 1.0e-03 |
| **ZV45724** | SRR6323866;NODE_45724_length_541_cov_3.835470 | 468 | HHRz III | 9.4e-07 | HHRz II | 1.8e-02 |
| **ZV28323** | SRR5864110;NODE_5877_length_1072_cov_3.852853 | 999 | TWRz | 9.7e-12 | TWRz | 9.7e-12 |

Nine of these zeta viruses contain delta ribozymes (DVRzs) in both polarity strands (Fig. 4). One zeta virus (ZV138865) contains type II HHRzs in both polarities, whereas four (ZV98882, ZV39584, ZV63649, and ZV45724) contain a type II HHRz in one polarity and a type III HHRz in the other (Fig. 4). In addition, one zeta virus (ZV130856) harbours a twister ribozyme (TWRz) in one polarity strand and a hairpin ribozyme (HPRz) in the opposite polarity, whereas another one (ZV5877) contains twister ribozymes in both polarities (Fig. 4). Interestingly, the TWRz identified in one polarity of ZV130856 displays an additional helix–loop element that is absent in the canonical TWRz structure, with the corresponding nucleotides highlighted in red in Fig. 4. Moreover, ZV5877, which contains two TWRzs, has a genome length of 999 nt, making it longer than all other zeta viruses reported so far.

AlphaFold 3 predictions of the tertiary structures of the putative endless proteins encoded in both polarity of these newly identified zeta viruses yielded pTM scores lower than 0.5 (data not shown), indicating low confidence in the resulting structural models.

Pairwise sequence identity was highly variable and generally low among the zeta viruses identified in this study, both at the nucleotide level (Fig. S2A) and among the proteins potentially encoded by ORF A and ORF B (Fig. S2B). Nucleotide sequence identities ranged from 28% to 58%, except for ZV52073 and ZV110734, both containing two delta ribozymes, which share 90% sequence identity. Amino acid sequence identities reached a maximum of 45%, except for six pairwise comparisons involving both proteins of zeta viruses containing delta ribozymes (ZV31793 vs ZV39444, ZV12952 vs ZV42858, and ZV52073 vs ZV110734) that showed amino acid identity ranging from 82 to 95%.

The newly identified zeta viruses were predominantly associated with environmental metatranscriptomic datasets, including sediments, microbial mats, wetlands, soils, freshwater environments, and compost. The only exception was ZV36159, containing DVRzs in both polarities, which was identified in a dataset of viral genomes associated with invertebrate samples, specifically *Frankliniella intonsa*.

### Identification and characterization of viroid-like RNAs with novel properties

In the libraries C30, CD95 and CD99, four additional contigs were selected among those corresponding to putative viroid-like RNAs (109, 17 and 31 contigs, respectively, for further characterization due to their interesting putative coding properties. Namely, the NODE_786 (784 nt length, coverage 1.76) in the CD95 library, NODE_1793 (721 nt length, coverage 2.12) in CD99 library, NODE_270217 (525 nt length, coverage 2.29) and NODE_184674 (608 nt length, coverage 9.71) in the C30 library, which corresponded to putative circular RNAs of 664, 685, 514, and 481 nt, respectively. BLASTn analyses did not show any significant identity between these contigs and previously reported RNA sequences in databases. All these RNA elements fold into rod-like or quasi rod-like secondary structures (Fig. 5A). INFERNAL analysis identified in these putative viroid-like RNAs a type III HHRz in each polarity strand (Fig. 5B). In vdlRNAs from CD95 and CD99 libraries, one of the two ribozymes displayed a mutation in the conserved catalytic core, where the canonical CUGA is replaced by CCGA (Fig. 5B). This mutation was already observed in some zeta viruses identified in this study.

**Fig. 5.**
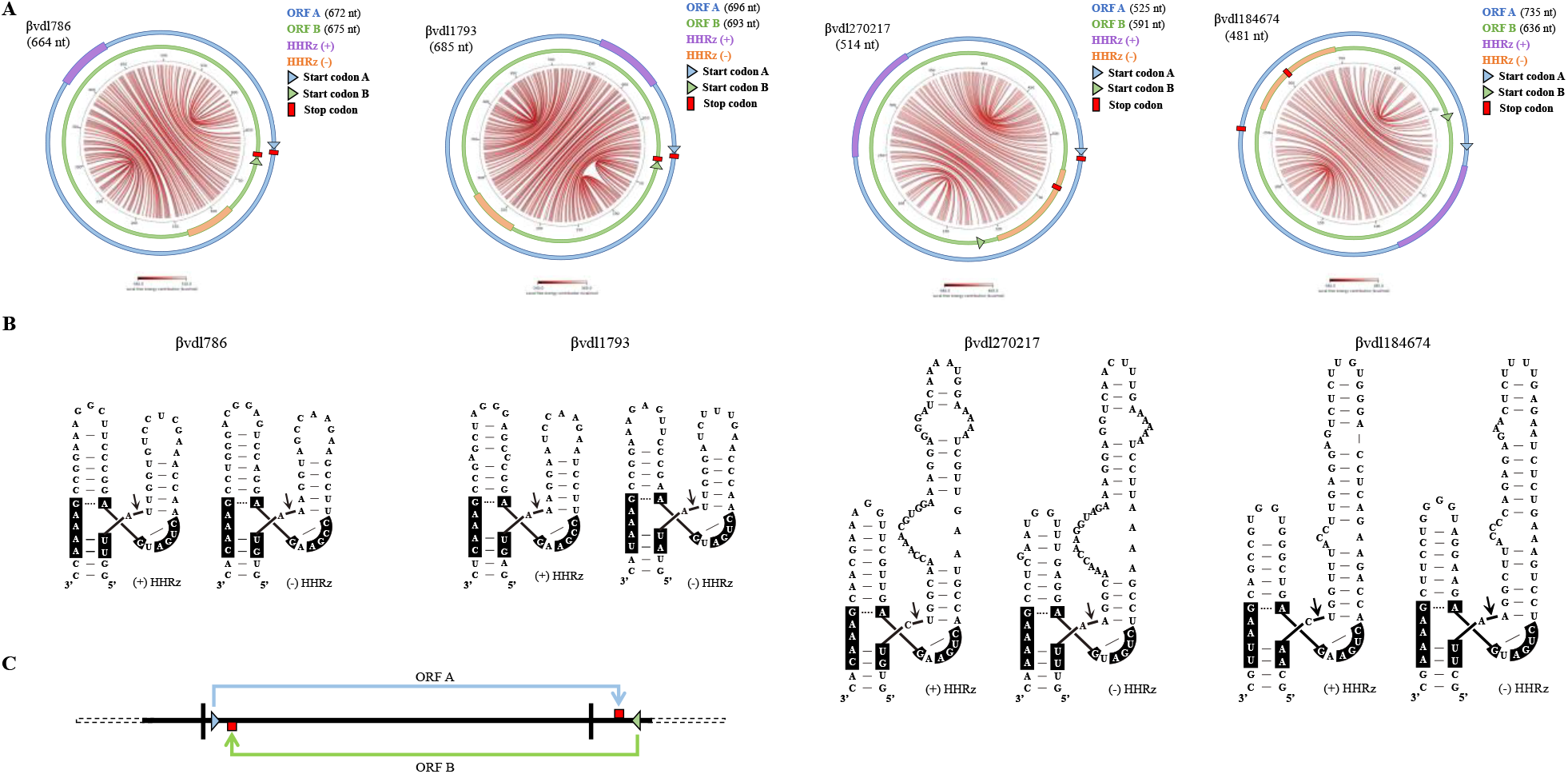
Structural features of betaviroid-like RNAs (βvdl786, βvdl1793, βvdl270217, and βvdl184674). (A) Circular plot representation of betaviroid-like RNA secondary structures of minimum free energy (see legend Figure 1 for more details). ORFs A and B (in blue and green, respectively), encoded by (+) and (-) polarity, respectively, are longer than the genomic RNA, with the start and stop codons indicated with a triangle and a square (red), respectively. The relative positions of the hammerhead ribozyme in (+) and (-) polarity strands are denoted in purple and orange, respectively. (B) Predicted secondary structure of hammerhead ribozymes in (+) (on the left) and (-) (on the right) polarity strands. (C) Schematic representation of a fragment of a head-to-tail multimeric betaviroid-like RNA.

The four viroid-like RNAs share a nucleotide sequence identity ranging from 47.95 to 56.97% with each other and exhibit similar structural and coding features. In both polarity strands, each RNA contains a single ORF that exceeds the length of the genomic RNA, with the stop codon falling within the second monomer of a multimeric RNA (Fig. 5C). This happens because the number of nucleotides composing the genome of these vdlRNAs is not a multiple of three (664, 685, 514, and 481 nt), as instead it happens in the case of zeta viruses. This scenario would imply the involvement of circular RNAs or, alternatively, of multimeric linear RNAs, likely generated during replication, as templates for translation. Additional studies are needed to further discriminate between these two possibilities.

Taking into consideration the presence of ORFs longer than the genomic RNA in both polarity strands, the name betaviroid-like RNAs has been proposed for these vdlRNAs to distinguish them from zeta viruses, that instead contain ORFs potentially generating endless tandem protein repeats. Therefore, the novel vdlRNAs identified in CD95, CD99 and C30 soil libraries were named betaviroid-like RNA 786 (βvdl786), betaviroid-like RNA 1793 (βvdl1793), betaviroid-like RNA 270217 (βvdl270217), and betaviroid-like RNA 184674 (βvdl184674).

BLASTp searches, revealed no significant similarity between the putative proteins encoded by the betaviroid-like RNAs and previously reported proteins. Pairwise comparison showed low amino acid sequence identity among the eight betaviroid-like encoded proteins, ranging from 9.77% to 33.33% amino acid identity. AlphaFold 3 predictions of their tertiary structures yielded low pTM scores (0.16-0.23; Fig. S3), indicating low confidence in the resulting structural models.

No polymorphic positions were identified by mapping the sequenced reads onto the respective βvdl786, βvdl1793, βvdl270217, and βvdl184674 genomic RNAs, supporting a strict conservation of genomic RNA sequences.

Among the four betaviroid-like RNAs identified, βvdl786 and βvdl1793 were selected for further investigation and detailed molecular characterization. The same experimental approach used for the zeta virus was applied to confirm the existence, the absence of a DNA counterpart and the ribozyme self-cleavage activity of the selected betaviroid-like RNAs. Reverse transcription, followed by PCR amplification with adjacent and complementary specific primers (Table S1) generated amplicons of the expected size for βvdl786 and βvdl1793 from soil samples CD95 and CD99, respectively (Fig. 6A). Cloning and Sanger sequencing of these amplicons confirmed the sequences obtained by RNA-seq analysis (βvdl786 and βvdl1793 GeneBank accession numbers: PZ584208 and PZ584209, respectively).

**Fig. 6.**
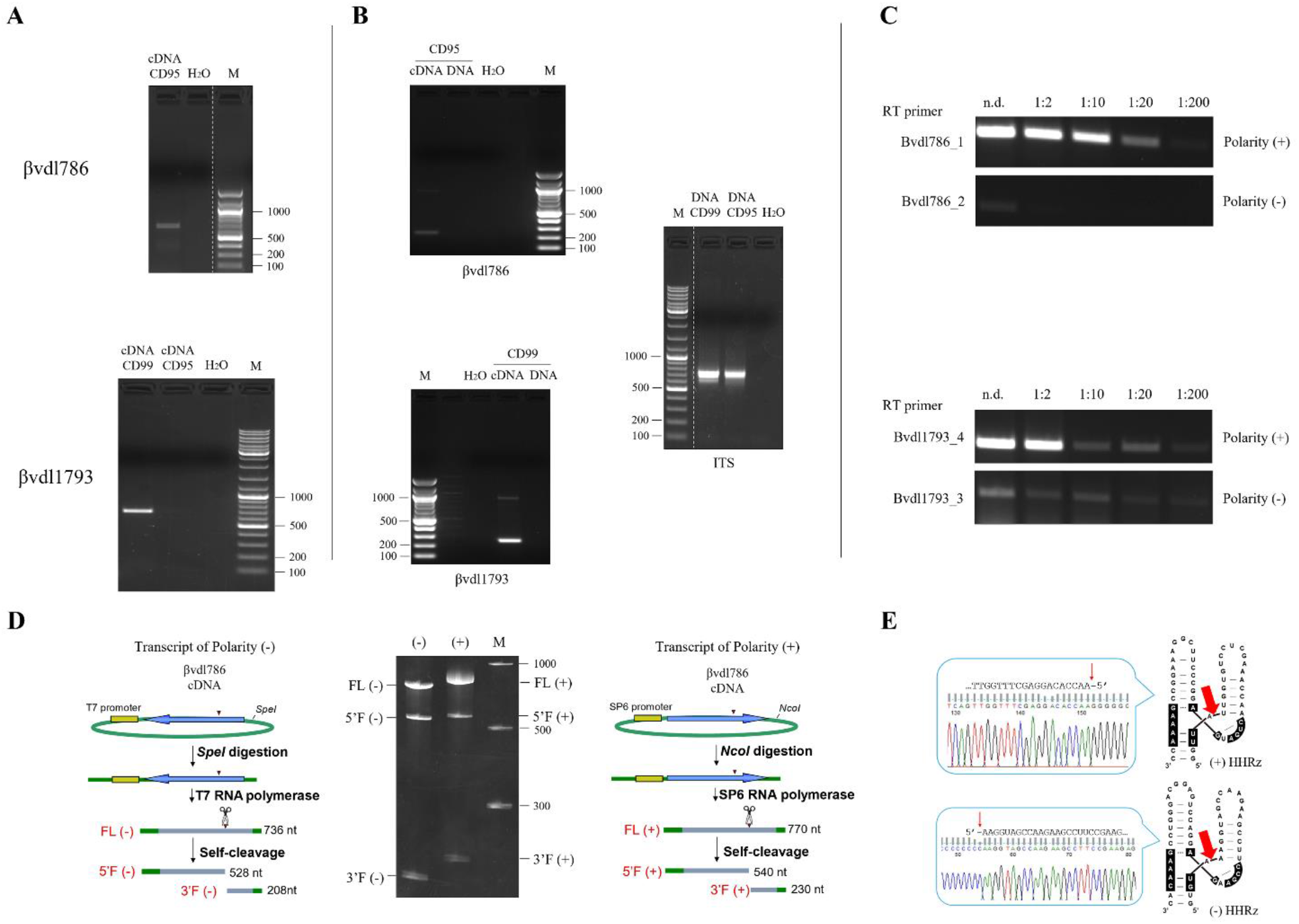
Molecular characterization of βvdl786 and βvdl1793. (A) RT-PCR amplification of full-length βvdl786 (664 nt, upper panel) and βvdl1793 (685 nt, bottom panel) from soil RNA samples CD95 and CD99, respectively, by using adjacent specific primers of opposite polarity (B) PCR amplification of a partial fragment of βvdl786 (248 nt, left upper panel) and βvdl1793 (256 nt, left bottom panel) using as template either cDNA or DNA preparations, to confirm the absence of DNA counterpart. In the right panel is shown the PCR amplification of CD95 and CD99 DNA with fungal universal primers ITS4/ITS5, as positive control (C) Strand-specific RT semi-quantitative PCR amplification for the determination of the βvdl786 (top panel) and βvdl1793 (bottom panel) most abundant polarity strand, denoted as (+) polarity by convention. n.d., not diluted. M, Gene ruler DNA ladder mix (Thermo Scientific). (D) Analysis of self-cleavage activity of βvdl786. Schematic representations of the plasmid and the βvdl786 RNA products of plus and minus polarity generated by *in vitro* transcription are shown on the right and left panels, respectively. Plasmid containing the monomeric cDNA sequence of βvdl786 was linearized and transcribed to produce full length monomeric transcripts (FL) of (+) (on the right) and (−) (on the left) polarity strands and the respective 5′ (5′F) and 3′ (3′F) fragments derived from the HHRz self-cleaving activity. Plasmid vector sequences are depicted in green, while in yellow and blue indicate the polymerase promoter and βvdl786 sequences, respectively. The scissors mark the position of the self-cleavage sites. The expected size of each RNA fragment is reported on the right. The middle panel shows the 5% PAGE analysis of the *in vitro* transcription of (+) and (-) βvdl786 monomers. M, RNA marker (Low range ssRNA ladder, New England Biolabs), with sizes (nt) indicated on the left. (E) The self-cleavage sites (red arrow) of the (+) and (-) HHRz (on the right) were determined by 5’ RACE of the 3’F fragments resulting from HHRz mediated cleavage of βvdl786 monomeric transcripts. The sequencing electropherograms of 5′ RACE products are shown on left site of the corresponding ribozyme.

To test whether the detected RNA sequences were also present in DNA form, DNA preparations from CD95 and CD99 soil samples were assayed by PCR using betaviroid-like specific primers amplifying sequence fragments of 248 and 256 nt for βvdl786 and βvdl1793, respectively (Table S1). cDNA from the same soil samples and fungal universal primers ITS4/ITS5 (Table S1) were used as positive controls (Fig. 6B). No βvdl786 and βvdl179 PCR amplificons were obtained from the DNA preparations, supporting the absence of DNA counterparts in the tested soils, while PCR amplification products of the expected size were obtained from the positive controls.

To assess the presence of RNAs of two polarity strands and to distinguish the (+) from the (-) strands based on their relative abundance, semiquantitative RT-PCR was performed using βvdl786 and βvdl1793 strand-specific primers (Table S1). Primers specific for (+) and (-) polarities were separately used for the strand-specific retro-transcription reaction, and several dilutions (not diluted, 1:2, 1:10, 1:20, 1:200) of the resulting cDNAs were amplified by PCR using specific primers for βvdl786 and βvdl1793 fragments. Both polarity strands were detected for both betaviroid-like RNAs. One RNA polarity of both βvdl786 and βvdl1793 accumulated to substantial higher steady state levels. Based on this evidence, this RNA polarity has been considered as the (+) one (Fig. 6C).

The self-cleavage activity of hammerhead ribozymes was assessed by *in vitro* transcription using plasmids containing the full-length cDNA monomer of βvdl786 as template. For both polarity strands, three bands corresponding to the full-length transcript and the two expected cleavage products were detected (Fig. 6D), indicating that the HHRz of both polarity strands are active *in vitro*. Finally, the precise self-cleavage sites in both polarity strands were confirmed by 5’ RACE analysis, which showed that the 3’ fragments generated by ribozyme activity possessed the expected 5’ termini (Fig. 6E). Altogether these data strongly support the involvement of a symmetric rolling circle mechanism in the replication of betaviroid-like RNAs.

### Identification of eleven novel betaviroid-like RNAs in publicly available datasets

To further investigate the global distribution of the betaviroid-like RNAs, we screened publicly available datasets. Eleven putative additional vdlRNAs, with genomes ranging from 286 to 685 nt in length and sharing similar structural and coding features to betaviroid-like RNAs, were identified in the dataset of viroid-like RNAs previously identified *in silico* by Forgia et al. (10) from metatranscriptomic data (Table 2). All these sequences were from environmental samples, including wetlands, stagnant water areas, plant litter, and boreal forest soils. Interestingly, the position of the stop codon in the second monomer is variable: in some betaviroid-like RNAs, it occurs just a few nucleotides from the beginning of the second monomer, as in the betaviroid-like RNAs characterized in this study, whereas in other cases, it is much distant, up to 260 nt (Table 2).

**Table 2.** List of putative betaviroid-like RNAs identified in metatranscriptomic datasets.

| ID | Contig | Genome size (nt) | Putative protein 1 (aa) | ORF A end position in the 2<sup>nd</sup> monomer (nt) | Putative protein 2 (aa) | ORF B end position in the 2<sup>nd</sup> monomer (nt) |
| --- | --- | --- | --- | --- | --- | --- |
| **βvdl786** | βvdl786 | 664 | 223 | 8 | 224 | 10 |
| **βvdl1793** | βvdl1793 | 685 | 230 | 8 | 231 | 11 |
| **βvdl270217** | βvdl270217 | 514 | 174 | 11 | 196 | 77 |
| **βvdl184674** | βvdl184674 | 481 | 244 | 254 | 211 | 155 |
| **βvdl79624** | SRR5247040;NODE_79624 | 676 | 228 | 11 | 231 | 20 |
| **βvdl15423** | SRR7080221;NODE_15423 | 658 | 224 | 17 | 224 | 17 |
| **βvdl17744** | SRR7006911;NODE_17744 | 541 | 184 | 14 | 218 | 116 |
| **βvdl20433** | SRR7004410;NODE_20433 | 676 | 295 | 212 | 275 | 152 |
| **βvdl41196** | SRR9016288;NODE_41196 | 286 | 106 | 35 | 181 | 260 |
| **βvdl34283** | SRR5829939;NODE_34283 | 523 | 182 | 26 | 178 | 14 |
| **βvdl40399** | SRR5829939;NODE_40399 | 478 | 243 | 254 | 163 | 14 |
| **βvdl590361** | SRR8535407;NODE_590361 | 458 | 170 | 55 | 157 | 16 |
| **βvdl34831** | ERR7672938_34831 | 655 | 221 | 8 | 220 | 8 |
| **βvdl37279** | SRR12926475_37279 | 619 | 230 | 71 | 209 | 8 |
| **βvdl725174** | SRR7080226_725174 | 637 | 218 | 17 | 215 | 8 |

No significant similarity of the novel circular betaviroid-like RNAs with any sequence previously deposited in databases was found by BLASTn analysis. Pairwise nucleotide sequence identities among the identified betaviroid-like RNAs ranged from 33.33% to 77.09% (Fig. S4A). Pairwise amino acid sequence identities among the predicted proteins encoded by the identified betaviroid-like RNAs reached a maximum of 59.55% (Fig. S4B). All the new identified betaviroid-like RNAs fold into rod-like secondary structures, sometimes slightly branched at one end, as predicted by RNAfold (Fig. 7).

**Fig. 7.**
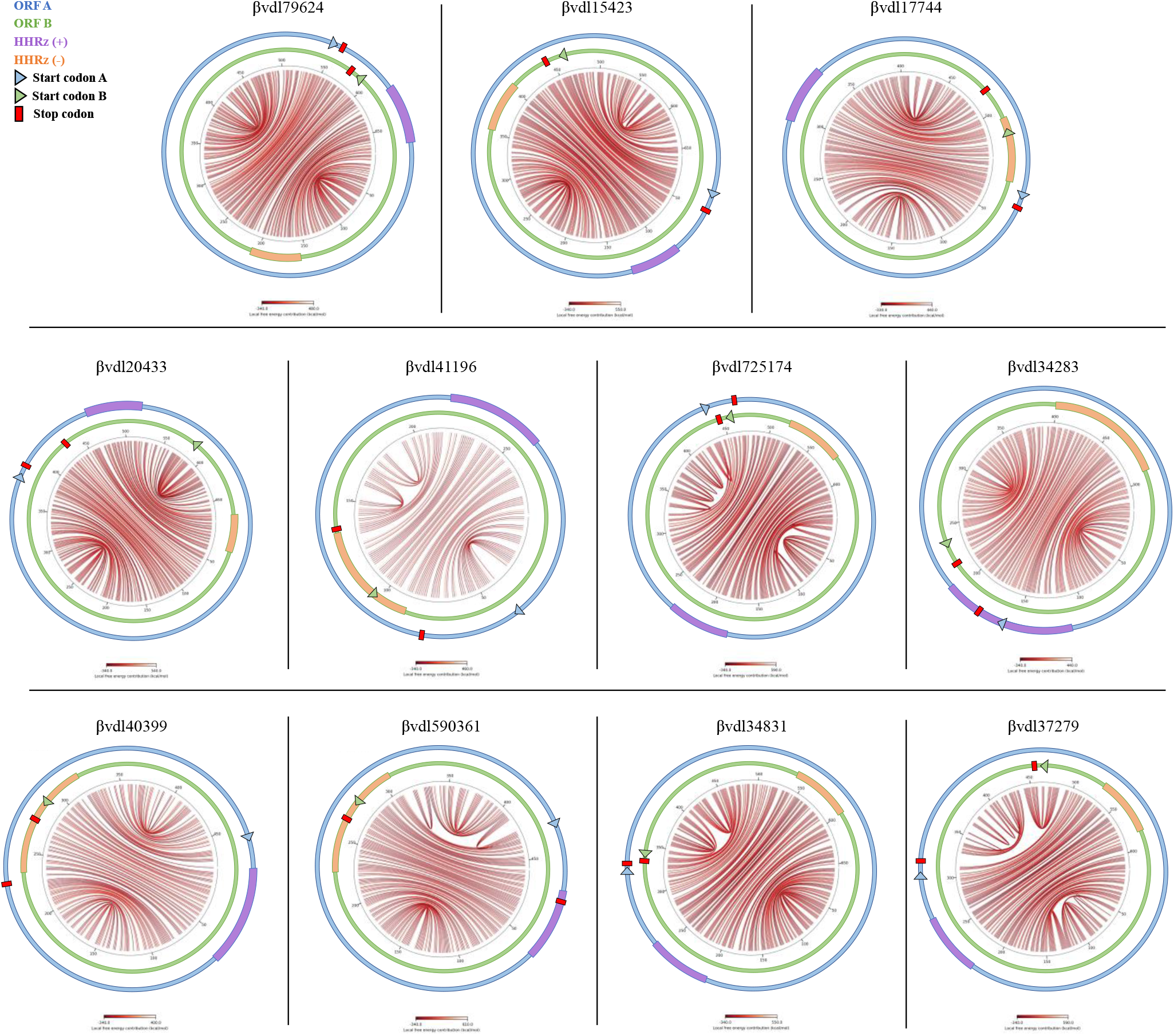
Circular plot of predicted secondary structures of betaviroid-like RNAs identified in metatranscriptomic datasets. The relative positions of ORFs A and B (blue and green), start (triangles) and stop (red squares) codons, and predicted hammerhead ribozymes (purple and orange) in both polarity strands are indicated.

INFERNAL analysis identified ambisense HHRz type III motifs in all putative betaviroid-like RNAs (Fig. 8). Three of them (βvdl34831, βvdl37279, and βvdl725174) exhibited the same mutation in the conserved catalytic core (CCGA instead of CUGA) present in βvdl786 and βvdl1793 HHRzs. The start and stop codons of the ORFs, in some cases, overlap with ribozyme motifs (e.g., βvdl41196, βvdl34283, βvdl40399, βvdl590361, βvdl37279), while in βvdl15423 the start and stop codons of both ORFs are paired in the rod-like secondary structure. Three vdlRNAs (βvdl34283, βvdl40399, βvdl590361), contained in both polarity strands HHRbzs differing from those found in the other vdlRNAs because of a long stem I of approximately 42-55 nt. Predicted proteins encoded by the ORFs of the novel betaviroid-like RNAs exhibit variable folding, with a low template modelling scores (pTM < 0.5) predicted by AlphaFold 3 (results not shown).

**Fig. 8.**
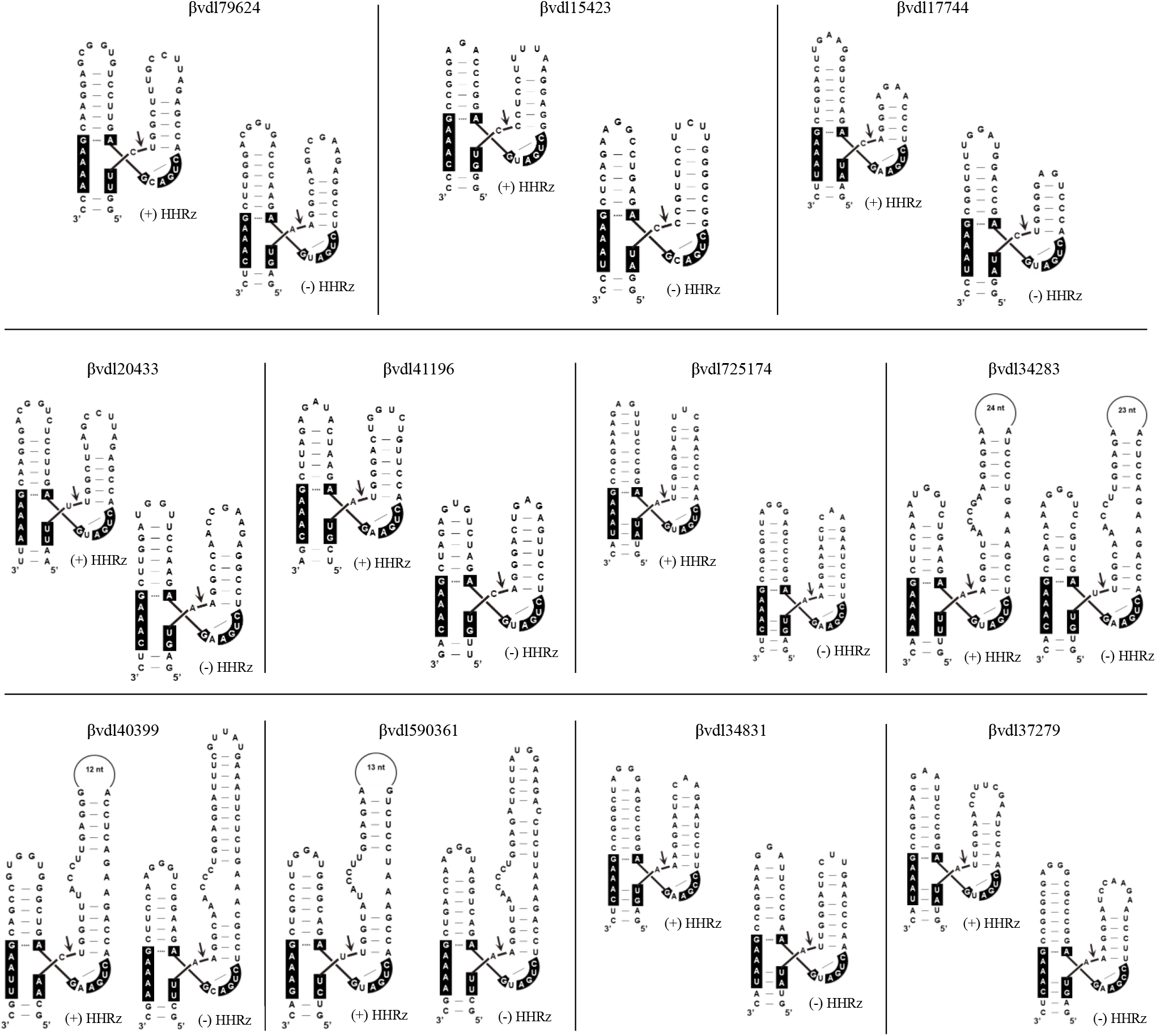
Secondary structure of (+) and (-) hammerhead ribozymes of betaviroid-like RNAs identified in metatranscriptomic datasets. The cleavage site is marked by an arrow and conserved nucleotides within the catalytic core are highlighted with a black background.

## DISCUSSION

VdlRNAs are a diverse group of infectious agents characterized by circular RNA genomes with highly base-paired secondary structures, often containing self-cleaving ribozymes and replicating through rolling-circle mechanisms. They include both non-coding and coding agents that differ in genome size, host range, and replication strategies, such as viroids, viroid-like satellite RNAs, deltaviruses, and other recently identified circular RNAs from fungi, bacteria, and environmental samples (10–14). Among these, zeta viruses, which encode putative endless proteins in each polarity strand, represent a recently proposed group identified only *in silico*, and whose existence had not yet been experimentally confirmed (15).

In the present study, a zeta virus identified in soil samples was molecularly characterized for the first time. The presence of (+) and (-) circular RNA forms, the absence of a DNA counterpart, and the self-cleavage activity of the hammerhead ribozymes in the two polarity strands were confirmed, providing evidence of replication through a symmetric rolling circle mechanism. We also identified and characterized a novel group of vdlRNAs, the betaviroid-like RNAs, which contain hammerhead ribozymes and ORFs slightly longer than their full-length circular genome, one in each polarity strand. The term betaviroid-like RNAs was adopted to highlight the viroid-like nature of these RNA elements and to distinguish them from conventional viruses, as no RNA-dependent RNA polymerase has been identified in these RNAs. Circularity, absence of a DNA form, and the catalytic activity of their hammerhead ribozymes were also validated for two of them. Although they share several features with zeta viruses, betaviroid-like RNAs exhibit a distinct genomic architecture. Notably, their genome is characterized by a nucleotide length that is never a multiple of three. This feature is consistent with a reading frame shift occurring after complete translation of a monomeric (linear or circular) genome, resulting in the eventual emergence of a termination codon. This situation contrasts sharply with zeta viruses, whose genomes consist of a number of nucleotides that is a multiple of three, allowing the translation of potentially endless proteins.

Both zeta viruses and betaviroid-like RNAs have been identified in environmental samples. Therefore, their host organisms remain still unknown. Nevertheless, this study provides compelling evidence of their existence in soil and offers experimental validation of their molecular characteristics. Moreover, the finding of twelve additional putative betaviroid-like RNAs in publicly available metatranscriptomic datasets further supports a broader distribution of this class of vdlRNA elements. In addition, the identification of zeta viruses containing ribozymes other than type III HHRzs (including DVRzs, TWRzs, HPRzs and type II HHrzs) further expands the known diversity of ribozymes associated with these RNA elements.

The unusual properties of the ORFs encoded by these vdlRNAs across both zeta viruses and betaviroid-like RNAs raise important questions regarding their translation strategies. One possibility is that translation occurs directly from the circular RNA through a reiterative translation cycle, the called rolling-circle translation. Such a mechanism has been demonstrated *in vitro*-synthesized circular RNA containing an endless ORF in bacterial (19) and eukaryotic cells (20). The rolling-circle translation mechanism has been also demonstrated to occur in endogenous circular RNAs such as the human circular-EGFR, a human circular RNA containing an endless or infinite ORF, the translation of which through a rolling circle and a programmed -1 ribosomal frameshifting inducing an out-of-frame stop codon produced a polymetric protein complex, the rolling translated EGFR (21). Different strategies of translation initiation have been proposed for circular RNAs, such as internal ribosomal entry site (IRES), N6-methyladenosine (m6A) elements, A-to-I RNA editing, and exon junction complex (EJC)-mediated initiation, as well as ribosome shunting, and repeat-associated non-AUG (RAN) translation (22). Nevertheless, the absence of strongly conserved canonical translation-initiation signals would not necessarily preclude translation from circular templates, as circular RNAs have been shown to support protein synthesis in the absence of an IRES, 5’ cap, or poly(A) tail (20). Alternatively, translation could proceed from linear multimeric RNAs, potentially corresponding to replication intermediates of the vdlRNAs. While translation from a circular RNA template presents challenges such as ribosome recruitment, in the linear-template model ribozymes might play a role in initiating translation, as it has recently been proposed for certain viruses (23).

The identification of ORFs in zeta viruses and betaviroid-like RNAs further expands the diversity of known viroid-like RNAs. In this respect, they show similarities to other circular RNAs, such as HDV and hepatitis Delta-like agents (family *Kolmioviridae*) (24), ambiviruses (25), and obelisks (13), that combine coding potential with hallmark features of viroids (including circular genomes, self-cleaving ribozymes, and rolling-circle replication). Evidence for translation capacity was shown also for the virusoid associated with rice yellow mottle virus (scRYMV), the unique among viroids and viroid-like satellite RNAs with coding capacity (26). While members of the *Kolmioviridae* family, obelisks and scRYMV encode proteins in one polarity strand, ambiviruses show coding capability in both polarity strands, encoding a putative RNA-dependent RNA polymerase in one polarity and another putative conserved protein with unknown function in the other polarity strand (10). Interestingly, zeta viruses, and betaviroid-like RNAs resemble ambiviruses in this ambisense coding capability. However, the arrangement of the coding sequences in the two polarity strands differs substantially. In ambiviruses, the two conserved ORFs are located in distinct, non-overlapping genomic regions of the opposite polarity strands (25). In contrast, the ORFs of zeta viruses and betaviroid-like RNAs span the entire genome in both polarities, representing an extreme form of ORF overlap that, to our knowledge, has not previously been described in a replicating RNA or virus (27). In addition, the genome organization of zeta viruses and betaviroid-like RNAs represents an extreme form of genome compression, in which the same nucleotide sequence simultaneously contributes to coding information in both strands while contributing also to structural and functional RNA elements involved in the replication of the genomic RNA and possibly in its translation.

In RNA viruses, overlapping coding sequences increase hypersensitivity to mutation, because a single nucleotide change may affect more than one coding region This genomic organization has been associated with greater mutational robustness to contrast high mutation rates (27). In this respect, the genomic organization of zeta viruses and betaviroid-like RNAs would suggest the need of an extreme mutational robustness for these replicons, likely to face deleterious mutations that may compromise both RNA structural elements and the coding capability which are overlapping in these vdlRNAs. Consistent with these considerations is the very low genetic variability observed for both zeta viruses and betaviroid-like RNAs in this study. The surprising genomic organization of zeta viruses and betaviroid-like RNAs open also interesting questions on their origin and evolutionary strategies that underscores the remarkable evolutionary plasticity of vdlRNAs.

Overall, these findings expand the known diversity of protein-coding vdlRNAs and highlight the remarkable genomic information compression of zeta viruses and betaviroid-like RNAs, providing new perspectives on their evolution and biological roles.

## MATERIALS AND METHODS

### RNA and DNA extractions, and bioinformatic analysis

Soil samples originally collected in 2019 from an orchard located in the municipality of San Godenzo (Tuscany, Italy) and previously used for the characterization of fungal communities associated with chestnut (*Castanea sativa*) trees by Venice et al. (17), were analyzed in the present study. Each soil sample was composed of five top-soil cores (200 cm^3^ each, at least two meters from each other) collected and pooled from close areas in the same field. Six soil samples were used for the total RNA extraction using RNeasy PowerSoil Total RNA kit (QIAGEN, Hilden, Germany) following manufacturer’s instructions. The six RNA preparations were sent to Novogene (Cambridge, United Kingdom; https://www.novogene.com/) for Illumina TruSeq Stranded Total RNA Library construction with riboZero Plant ribosomal removal and subsequent high-throughput sequencing using the Illumina NovaSeq 6000 (2×150bp). The resulting RNAseq libraries were generated as part of a broader study aimed at characterizing the soil viromes, with the corresponding virome analysis being reported in a separate manuscript currently in preparation. The six RNAseq libraries were denoted C30, CD50, CD95, CD99, C105 and C110. Total DNA was extracted from the same soil samples with the FastDNA™ SPIN Kit for Soil (MP Biomedicals, Europe, https://www.mpbio.com/eu/) according to manufacturer’s instructions.

Quality control of raw reads was performed using the FASTX-Toolkit (https://github.com/agordon/fastx_toolkit). Reads were then pre-processed with cutadapt (28) to trim adapter sequences, discard low-quality and overly short reads (Phred score Q<20, post-trimming length L<50 bp), and remove undetermined nucleotides (N). Reads passing these filters were subjected to *de novo* assembly using SPAdes (v3.13.0) (29), applying a multi k-mer strategy (k-mer sizes from 51 to 121 bp, in increments of 10). Contigs were functionally annotated via a local BLASTx search against the NCBI non-redundant protein database (downloaded August 4, 2023), implemented with DIAMOND (v2.1.7) (30).

The resulted assembled contigs were analysed by INFERNAL (v1.1.2) algorithm (16) for the identification of RNAs containing ribozymes, using covariance models from Forgia et al. (10) and those built on ribozyme structures from RFAM database v14 (31). Sequences with paired ribozymes, circular topology (inferred by 5’ -3’ overlaps in the assembled contig) and no BLAST annotation were selected for further analysis. To assess sequence variability, raw sequencing reads were mapped to the reference vdlRNA contig of interest using Geneious mapper with medium–low sensitivity and default parameters unless otherwise specified (Geneious Prime v.2026.0, Biomatters Ltd.). Variant calling analysis was performed by setting a minimum coverage threshold of 50 reads, a minimum variant frequency of 0.3%, consistent with the reported Illumina sequencing error rate (32), a maximum variant *p*-value of 1 × 10⁻⁴, and a minimum strand-bias *p*-value of 0.01.

Nucleotide and amino acid sequence similarity analyses of zeta viruses and betaviroid-like RNAs, and their predicted encoded proteins, respectively, were performed using Clustal Omega through the EMBL-EBI Job Dispatcher web server (33). Pairwise identity matrices generated from the multiple sequence alignments were subsequently used to produce heatmaps in RStudio using ggplot2 package.

Searches for additional zeta viruses and betaviroid-like RNAs in publicly available metatranscriptomic datasets were performed using the circular sequences reported by Forgia et al. (10) as starting templates. Circular monomers shorter than 2,000 nt were computationally dimerized *in silico*, the generated dimeric sequences were translated and only sequences with ORF longer than the monomeric length in both polarity strands were retained. The selected sequences were then separated according to whether their genomic length was or was not a multiple of three, corresponding to the expected characteristics of putative zeta viruses and betaviroid-like RNAs, respectively. Subsequent ribozyme identification using Infernal was performed as described above. Multiple sequence alignments of the nucleotide sequences and of the predicted proteins were generated using Clustal Omega (33).

RNA secondary structures were predicted with RNAfold web server, ViennaRNA package 2.0 (34). RNAeval 2.6.3 ViennaRNA outputs were used to generate circular plots with Jupyter Notebook environment (35), in Python 3 environment (36) with libraries for data analysis and visualization (i.e., NumPy, plotly, matplotlib).

ORFs were searched using Translate tool (Expasy) (37). The tertiary structure predictions of the putative encoded proteins were obtained by AlphaFold Server powered by AlphaFold 3 (38). Because most endless ORFs of zeta viruses contained multiple AUG codons, the translation initiation site could not be unambiguously assigned. Therefore, for each polarity strand of zeta viruses, the start codon of the endless ORF yielding the protein with the highest predicted template modelling (pTM) score by AlphaFold 3 server was selected for subsequent structural and alignment analyses.

### RT-PCR and cloning

Total nucleic acids were retro-transcribed (RT) using random hexamers and Superscript IV reverse transcriptase (ThermoFisher Scientific, Waltham, MA, USA). Two microliters of the cDNA reaction served as template for PCR amplification using 0.25 units of GoTaq DNA Polymerase (Promega, Madison, WI, United States) in a 25 µL mix containing 1X Green GoTaq Reaction Buffer, 0.2 mM of dNTPs and 0.2 µM of each specific primer (Table S1) and using the following cycling conditions: initial denaturation at 94 °C for 3 min, followed by 32 cycles at 94 °C for 30 sec, 57 °C for 30 sec, 72 °C for 30 sec, and a final extension step at 72 °C for 5 min. Amplification products of the expected size were excised from the agarose gel, purified, cloned into a pGEM-T Easy vector (Promega, Madison, WI, United States) and sequenced by Sanger sequencing (Macrogen, Amsterdam, Netherlands).

Quantification of each polarity strand of vdlRNAs was assessed by strand-specific RT followed by semi-quantitative PCR amplification. For each polarity strand, a reverse transcription was performed using a strand specific primer, then serial cDNA dilutions (not diluted, 1:2, 1:10, 1:20, 1:200) served as template for a PCR amplification as described above using primers indicated in Table S1.

### Analyses of RNA self-cleavage

Monomeric transcripts of both polarity strands were obtained by *in vitro* transcription of pGEM-T Easy plasmids containing the full-length cDNA sequence of the vdlRNAs. Recombinant plasmids were linearized with the appropriate restriction enzyme, purified by phenol–chloroform extraction followed by ethanol precipitation and used as templates for *in vitro* transcription with T7 (ThermoFisher Scientific, Wattham, USA) or SP6 RNA polymerase (New England Biolabs, Ipswich, MA, USA). Transcription reactions were analysed on denaturing 5% PAGE (8 M urea, 1 X TBE: 89 mM Tris, 89 mM boric acid, 2.5 mM EDTA, pH 8.3), stained with ethidium bromide, and visualized under UV light. The 3′ self-cleavage products generated during the *in vitro* transcription were eluted from the acrylamide gels and their 5’ terminal sequence of RNA was determined by 5’ Rapid Amplification of cDNA Ends (RACE) analysis. Briefly, self-cleavage 3’ RNA fragments were excised from the PAGE, purified by phenol– chloroform extraction and ethanol precipitation, and reverse transcribed using specific primers (Table S1). After addition of a poly(G) tail by terminal deoxynucleotidyl transferase (Promega, Madison, WI, USA), the tailed cDNAs were PCR amplified using GoTaq DNA polymerase (Promega, Madison, WI, United States) with the primers polyC RACE and a specific primer, nested to that used for the cDNA synthesis (Table S1). PCR amplicons were gel-purified, cloned, and Sanger sequenced.

## DATA AVAILABILITY

The RNA-seq raw data generated in this study have been deposited in the NCBI Sequence Read Archive (SRA) under BioProject accession number PRJNA1489414.

## ACKNOWLEDGMENTS

This research was funded by the EU LIFE program in the framework of the LIFE MycoRestore project (LIFE18/CCA/ES/001110) and partially supported by the Microbes-4-Climate – Project No. 101131818 funded by the European Union’s Horizon Europe program.

Nadia Serale: Data curation, Formal analysis, Investigation, Methodology, Validation, Writing – original draft, Writing – review & editing. Michela Chiumenti: Data curation, Formal analysis, Methodology, Software, Writing – review & editing. Paolo Mussano: Investigation. Silvia Rotunno Investigation. Francesco Venice: Resources. Gianni Della Rocca: Resources. Massimo Turina: Writing – review & editing. Antonietta Mello: Funding acquisition, Resources, Writing – review & editing. Laura Miozzi: Funding acquisition, Writing – review & editing. Francesco Di Serio: Conceptualization, Supervision, Writing – original draft, Writing – review & editing. Beatriz Navarro: Conceptualization, Data curation, Formal analysis, Supervision, Writing – original draft, Writing – review & editing.

## SUPPLEMENTAL MATERIAL

**Table S1.**
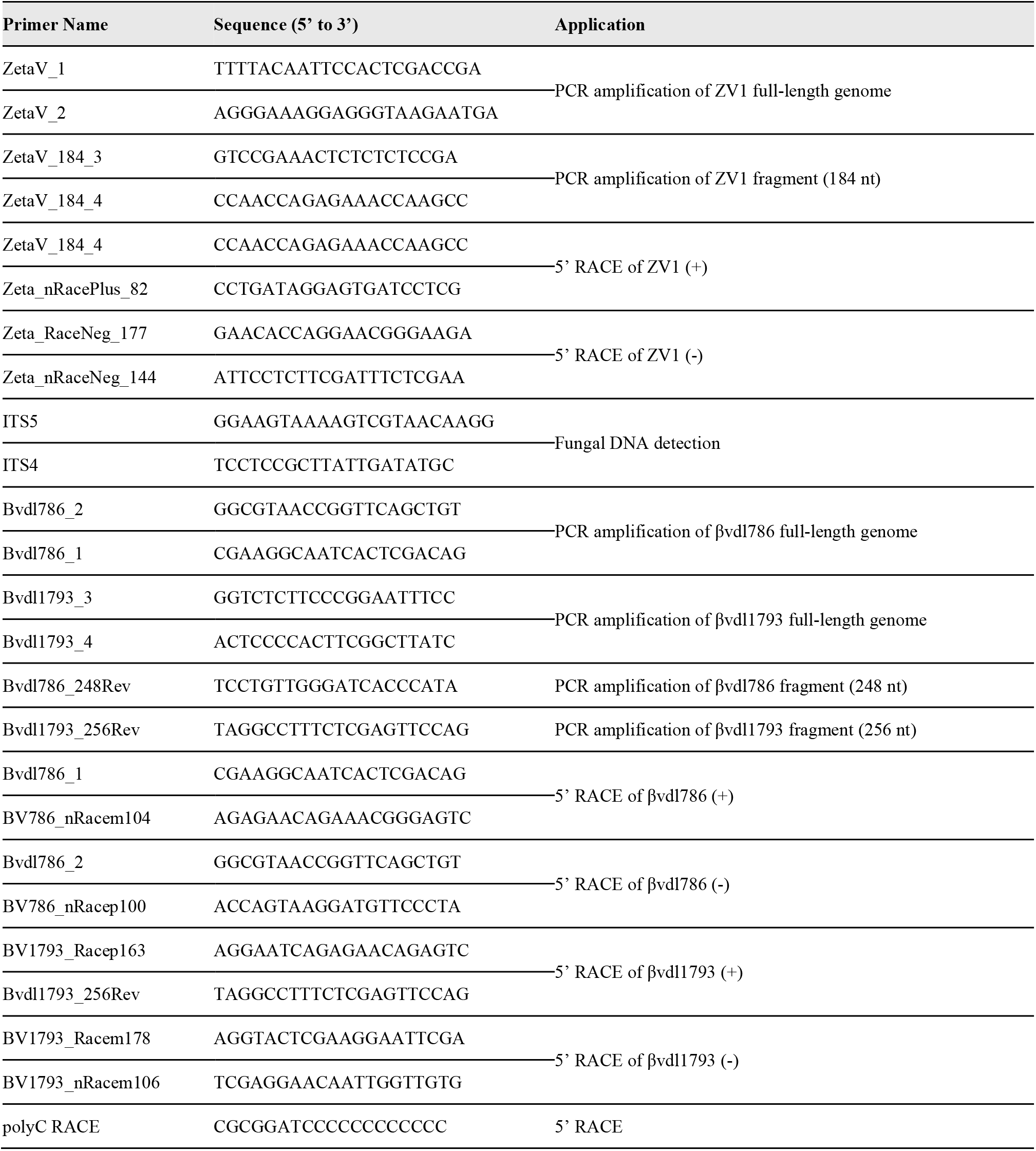
List of primers used in this study.

**Fig. S1.**
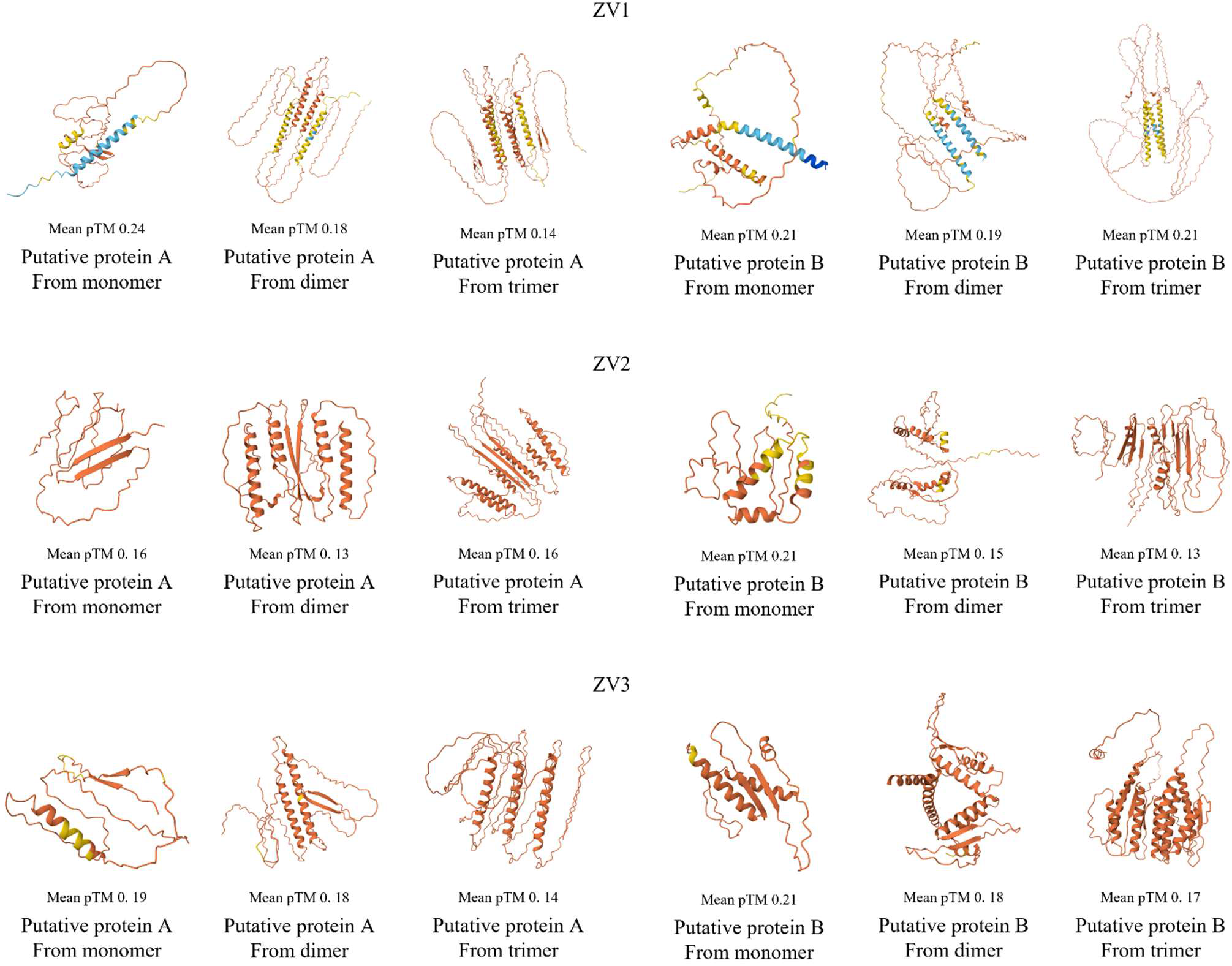
Tertiary structures of zeta virus ZV1, ZV2, and ZV3 putative proteins (A and B) predicted with AlphaFold Server powered by AlphaFold 3.

**Fig. S2.**
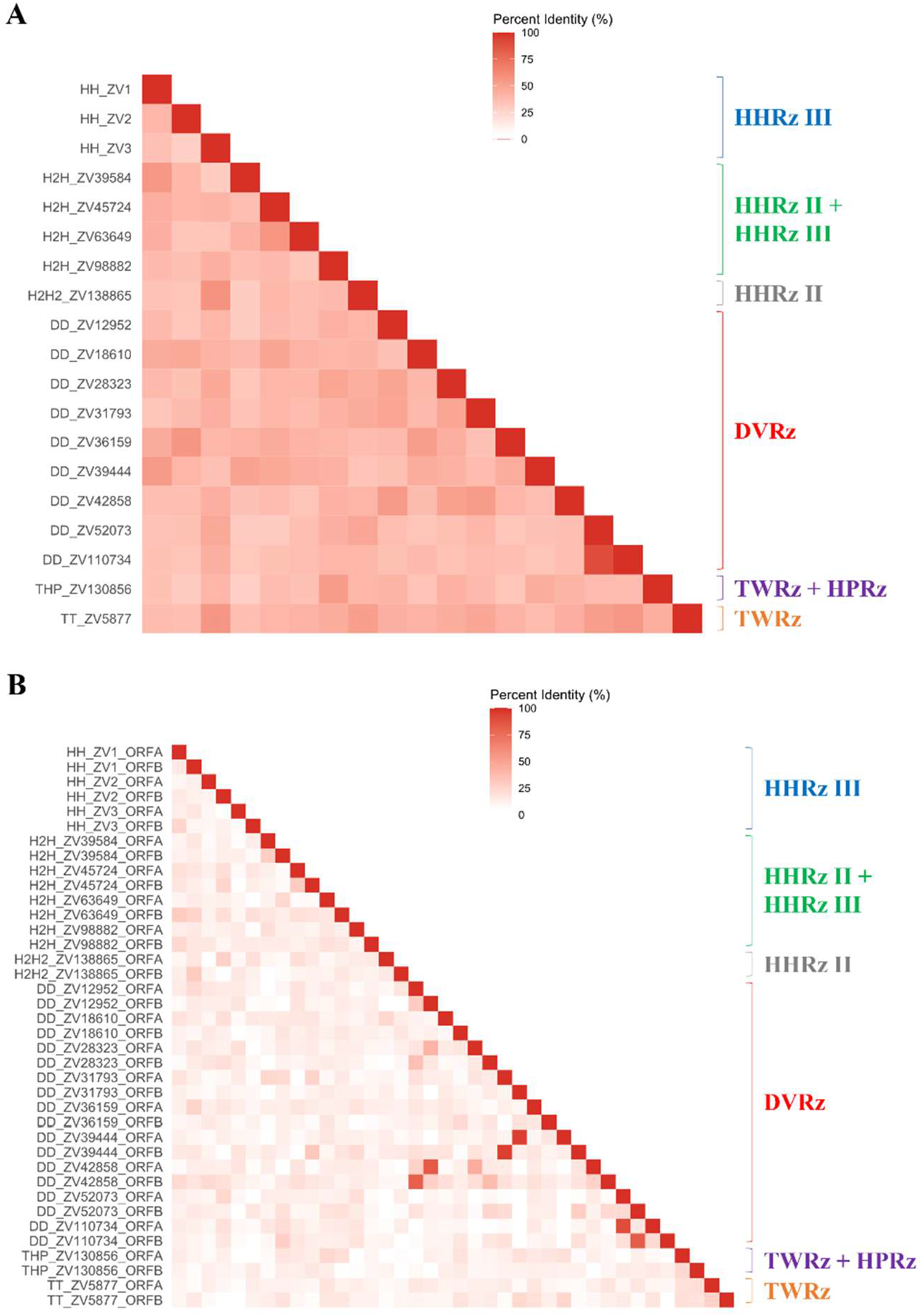
Nucleotide and amino acid sequence identities of zeta viruses. (A) Heatmap representing percent identity matrix of nucleotide sequence of all identified zeta viruses with different type of ribozymes (H: type III hammerhead; H2: type II hammerhead; D: delta; T: twister; HP: hairpin ribozymes). (B) Heatmap representing percent identity matrix of amino acid sequence of putative proteins (ORF A and ORF B) of zeta viruses. DVRz: delta ribozyme; II HHRz: type II hammerhead ribozymes; III HHRz: type III hammerhead ribozymes; HPRz: hairpin ribozyme; TWRz: twister ribozyme.

**Fig. S3.**
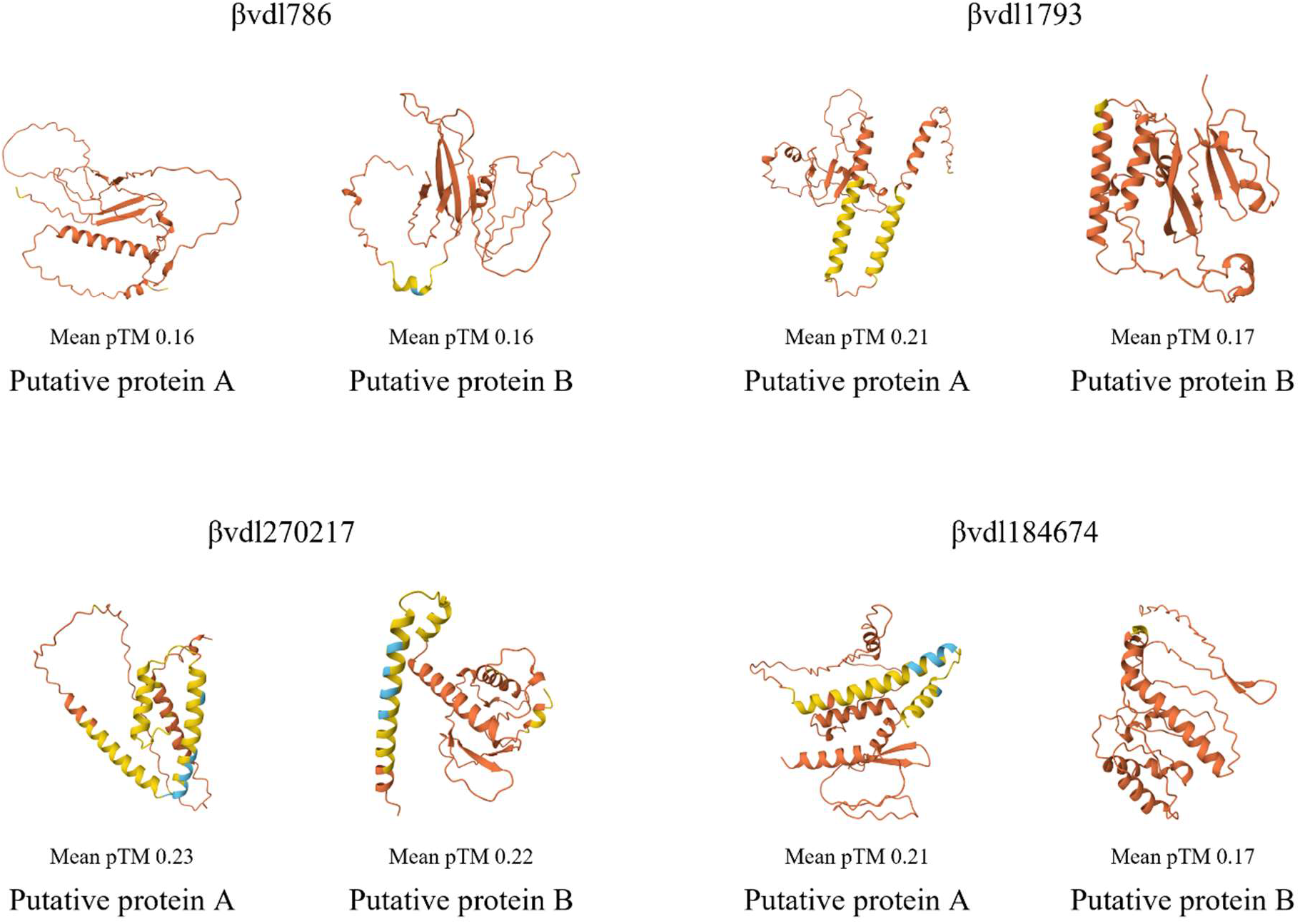
Tertiary structures of putative proteins encoded by betaviroid-like RNAs (βvdl786, βvdl1793, βvdl270217, and βvdl184674,) predicted with AlphaFold Server powered by AlphaFold 3.

**Fig. S4.**
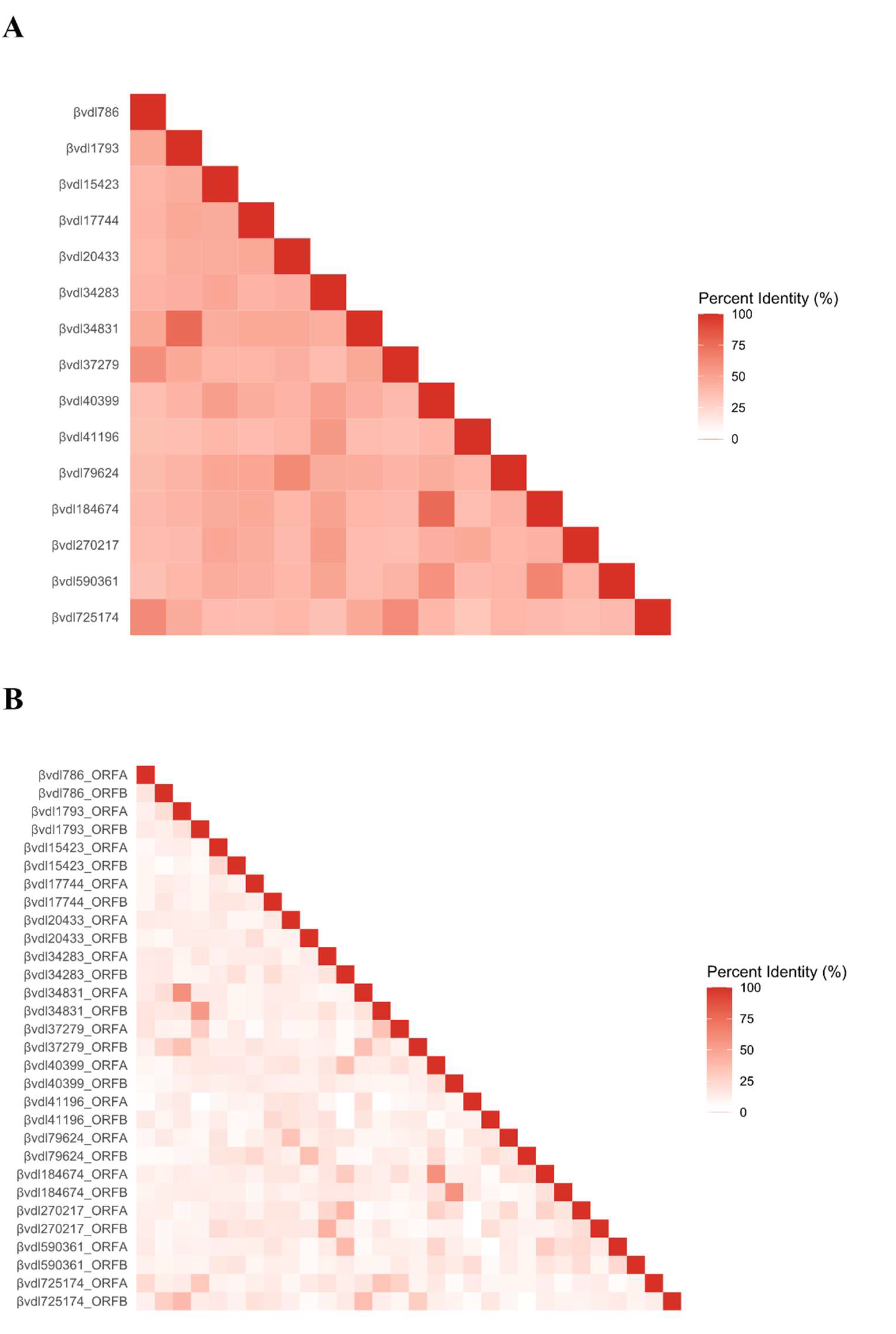
Nucleotide and amino acid sequence identities of betaviroid-like RNAs. (A) Heatmap representing percent identity matrix of all identified betaviroid-like RNA nucleotide sequences. (B) Heatmap representing percent identity matrix of amino acid sequences of putative proteins (ORF A and ORF B) encoded by the fifteen betaviroid-like RNAs.

